# Surveillance of Vermont wildlife in 2021-2022 reveals no detected SARS-CoV-2 viral RNA

**DOI:** 10.1101/2023.04.25.538264

**Authors:** Hannah W. Despres, Margaret G. Mills, Madaline M. Schmidt, Jolene Gov, Yael Perez, Mars Jindrich, Allison M. L. Crawford, Warren T. Kohl, Elias Rosenblatt, Hannah C. Kubinski, Benjamin C. Simmons, Miles C. Nippes, Anne J. Goldenberg, Kristina E. Murtha, Samantha Nicoloro, Mia J. Harris, Avery C. Feeley, Taylor K. Gelinas, Maeve K. Cronin, Robert S. Frederick, Matthew Thomas, Meaghan E. Johnson, James Murphy, Elle B. Lenzini, Peter A. Carr, Danielle H. Berger, Soham P. Mehta, Christopher J. Floreani, Amelia C. Koval, Aleah L. Young, Jess H. Fish, Jack Wallace, Ella Chaney, Grace Ushay, Rebecca S. Ross, Erin M. Vostal, Maya C. Thisner, Kyliegh E. Gonet, Owen C. Deane, Kari R. Pelletiere, Vegas C. Rockafeller, Madeline Waterman, Tyler W. Barry, Catriona C. Goering, Sarah D. Shipman, Allie C. Shiers, Claire E. Reilly, Alanna M. Duff, David J. Shirley, Keith R. Jerome, Ailyn C. Pérez-Osorio, Alexander L. Greninger, Nick Fortin, Brittany A. Mosher, Emily A. Bruce

## Abstract

Previous studies have documented natural infections of SARS-CoV-2 in various domestic and wild animals. More recently, studies have been published noting the susceptibility of members of the Cervidae family, and infections in both wild and captive cervid populations. In this study, we investigated the presence of SARS-CoV-2 in mammalian wildlife within the state of Vermont. 739 nasal or throat samples were collected from wildlife throughout the state during the 2021 and 2022 harvest season. Data was collected from red and gray foxes (*Vulpes vulples* and *Urocyon cineroargentus*, respectively), fishers (*Martes pennati*), river otters (*Lutra canadensis*), coyotes (*Canis lantrans*), bobcats (*Lynx rufus rufus*), black bears (*Ursus americanus*), and white-tailed deer (*Odocoileus virginianus*). Samples were tested for the presence of SARS-CoV-2 via quantitative RT-qPCR using the CDC N1/N2 primer set and/or the WHO-E gene primer set. Our results indicate that no sampled wildlife were positive for SARS-CoV-2. This finding is surprising, given that most published North America studies have found SARS-CoV-2 within their deer populations. The absence of SARS-CoV-2 RNA in populations sampled here may provide insights in to the various environmental and anthropogenic factors that reduce spillover and spread in North American’s wildlife populations.

## Introduction

Severe acute respiratory syndrome associated coronavirus-2 (SARS-CoV-2), the virus that causes COVID-19, is most recognized for its ability to easily transmit from person-to-person. Recently, natural infections in a range of domestic and wild animals have also been documented^1–4^. With every new animal infected, the zoonotic potential of SARS-CoV-2 increases. Animal species that facilitate within-species transmission of SARS-CoV-2 are possible new reservoirs of the virus, and this transmission could lead to evolutionary changes in the virus that would pose a risk to humans upon reintroduction. In fact, this exact scenario occurred during 2020, with SARS-CoV-2 infection documented in farmed minks ^5,6^. Notably, the Netherlands reported five different outbreak events in 2020, resulting in over 50% of mink farms having animals that tested positive for SARS-CoV-2. At over half of the farms with positive animals, employees also tested positive for SARS-CoV-2. Sequencing data from both the mink and humans suggests that both spillover, the transmission of disease from animals to humans, and spillback, the transmission of disease from humans to animals, occurred several times between these two populations^6^. Infected animals were detected in mink farms in multiple other countries which led to the selective culling of animals at affected farms, as well as the culling of all (>17 million) mink in Denmark, to reduce the risk of spillover^7,8^.

Spillover, the transmission of disease from animal to human, and spill back, the transmission of disease from animals back into people, are both thought to be relatively rare in Vermont (VT). However, multiple recent studies in North America have shown that members of the Cervidae family are susceptible to SARS-CoV-2. We hypothesized that SARS-CoV-2 might be circulating in Vermont deer and wildlife, given the numerous reports of infections within wild deer populations^4,9–15^, and laboratory infections showing vertical^16^ and horizontal transmission^17^. North American deer are of particular concern as they are common, interact with humans, and are also domestically farmed. All three of these factors create opportunities for spillover and spillback events. The 2021 estimate for Vermont’s white-tailed deer population was approximately 133,000^18^, which is about a 1:5 deer-to-person ratio within the state^19^. While SARS-CoV-2 has been detected in wildlife in several US states and Canadian provinces, there is currently no published data on the virus in wildlife in the state of Vermont.

In this study, we examined the prevalence of SARS-CoV-2 viral RNA in a variety of animals native to Vermont via reverse transcription quantitative polymerase-chain reaction (RT-qPCR), using two different primer sets specific to SARS-CoV-2. We sampled fur-bearing animals including red and grey foxes (*Vulpes vulples* and *Urocyon cineroargentus*, respectively), fishers (*Martes pennati*), otters (*Lutra canadensis*), coyotes (*Canis lantrans*), bobcats (*Lynx rufus rufus*), and big-game animals including white-tailed deer (*Odocoileus virginianus*) and black bears (*Ursus americanus*) over the 2021 and 2022 hunting seasons.

## Results

Our SARS-CoV-2 surveillance effort covered the state of Vermont through the hunting and trapping seasons of 2021 (Oct 2021-March 2022) and the hunting season of 2022 (Oct-Nov 2022). We prioritized white-tailed deer, given their prevalence, potential interaction with humans, and our ability to collect high-quality samples for processing. In addition to white-tailed deer, the 2021 season also included a variety of fur-bearing animals that are commonly trapped in VT, including foxes, fishers, otters, coyotes, and bobcats. In 2021, we sampled 17 white-tailed deer as well as 250 fur-bearers (Table 1). However, most of our white-tailed deer sampling occurred during the 2022 season, where we were able to sample 470 white-tailed deer as well as 2 black bears (Table 1). Sampled animals were harvested across the state of Vermont, generating samples from a broad geographic range (Figure 1, Figure S1). At the conclusion of the 2021 season, we extracted RNA from all collected samples and performed RT-qPCR using the Centers for Disease Control and Prevention (CDC) SARS-CoV-2 N1 and N2 primer set for the structural nucleocapsid protein^20^ to test for the presence of viral RNA. We found no detectable SARS-CoV-2 RNA within any sample from the 2021 season (n = 272). Positive control wells on each plate amplified as expected, as did the internal control included in each sample extraction to confirm RNA integrity and rule out PCR inhibition. (Dataset S1).

**Table 1.** Number of samples collected by species type for each season.

| Species | 2021 Season | 2022 Season |
| --- | --- | --- |
| **White-tailed Deer**<br>(*Odocoileus virginianus*) | 17 | 470 |
| **Foxes**<br>(*Vulpes vulpes* & *Urocyon cinereoargenteus*) | 19 | 0 |
| **Fishers**<br>(*Martes pennati*) | 77 | 0 |
| **Otters**<br>(*Lutra canadensis*) | 43 | 0 |
| **Coyotes**<br>(*Canis latrans*) | 32 | 0 |
| **Bobcats**<br>(*Lynx rufus rufus*) | 79 | 0 |
| **Black bears**<br>(*Ursus americanus*) | 0 | 2 |

**Figure 1.**
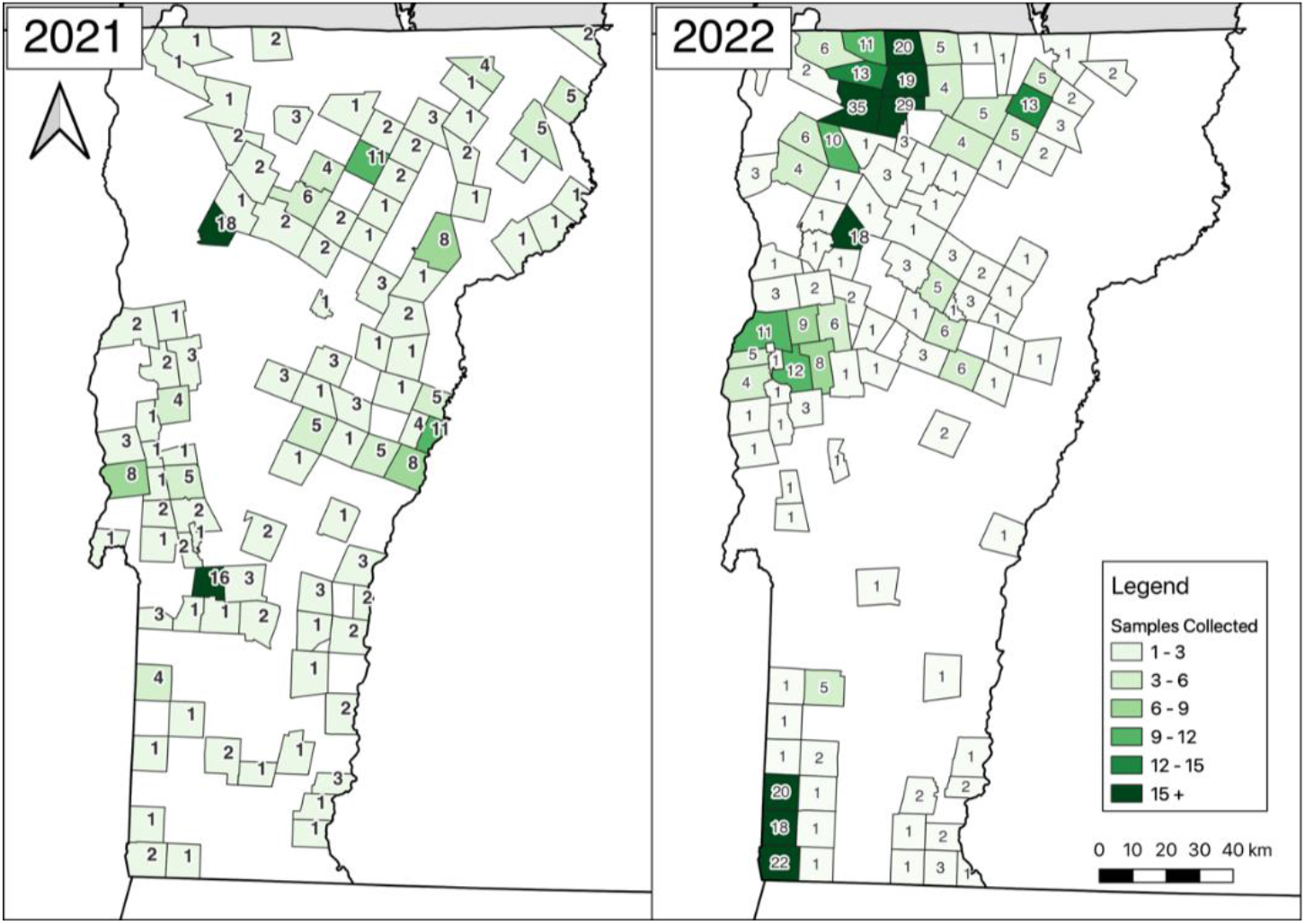
Geographic distribution of specimen harvest. Geographic distribution of wildlife sampled for SARS-CoV-2 in Vermont during the 2021 and 2022 hunting seasons is shown. Specimens are shown based on the reported town where the harvest occurred and colored according to the number of samples collected from each location. Graphs were generated using QGIS.

At the conclusion of the deer sampling period in 2022, we thawed all collected samples, divided each sample into two aliquots (one for RT-qPCR and one for future viral isolation), re-froze samples at -80°C, and performed RNA extraction and RT-qPCR as above for the 2021 season samples. Surprisingly, when we analyzed samples from the 2022 season (n = 472), we observed a positivity rate of 28.2% of samples positive for both the N1 and N2 primers (133/472). The average cycle threshold (C_T_) for these samples was 36.6 for N1 and 38.0 for N2 (SD = 1.3 and 1.4, respectively). There were multiple additional samples positive by either the N1 or N2 primer sets, but not by both (N1 only = 28 samples, N2 only = 56 samples) (Dataset S1). The suddenly high number of positive samples, paired with the high average C_T_ values and the lack of any samples with a C_T_<30 for N1 or C_T_<33 for N2 raised concerns that these initial numbers from the 2022 season may have been the result of contamination. In the period between processing the 2021 and 2022 samples, The University of Vermont’s laboratory began a separate project that involved *in vitro* expression of the SARS-CoV-2 nucleocapsid protein, and thus a DNA construct containing the sequences recognized by the N1 and N2 primer sets was newly present in the general laboratory environment.

Therefore, we set out to determine if the positive results seen with the N1/N2 primers were authentic or the result of plasmid DNA contamination in the laboratory environment from the University of Vermont during the sample aliquoting, before any sample analysis. First, we performed environmental swabbing of commonly used items and surfaces within the laboratory, including within the biosafety cabinet used to aliquot the wildlife specimens before RT-qPCR testing, the pipettes used for aliquoting, the laboratory bench, and pipettes. We detected SARS-CoV-2 N nucleic acids on all surface swabs with both the N1 and N2 primers with C_T_s as low as 23.6 (Dataset S2). None of the negative controls for the RT-qPCR reaction amplified. To determine if we were detecting RNA or DNA contamination, we next performed a quantitative polymerase chain reaction (qPCR) in which the typical incubation for reverse transcription was omitted, instead beginning directly with a 95° C step to deactivate RT and activate hot-start Taq. The positive controls (remnant SARS-CoV-2 positive clinical specimen) included in these experiments exhibited an average N1 C_T_ 5.4±0.6 cycles higher in qPCR experiment than in RT-qPCR, as expected for samples where the input material was RNA rather than DNA (Figure 2). Two of the three positive controls were undetectable with the N2 primer set in qPCR experiments; for the third, the N2 C_T_ was 1.8 cycles higher in qPCR experiment than in RT-qPCR. In contrast, all laboratory sites sampled (except the biosafety cabinet floor, which had the highest C_T_ originally) showed consistent C_T_s between RT-qPCR and qPCR reactions (average N1 C_T_ 0.3±0.9 cycles higher in qPCR experiment than in RT-qPCR), suggesting that the surface contamination consisted of DNA rather than RNA (Figure 2).

**Figure 2.**
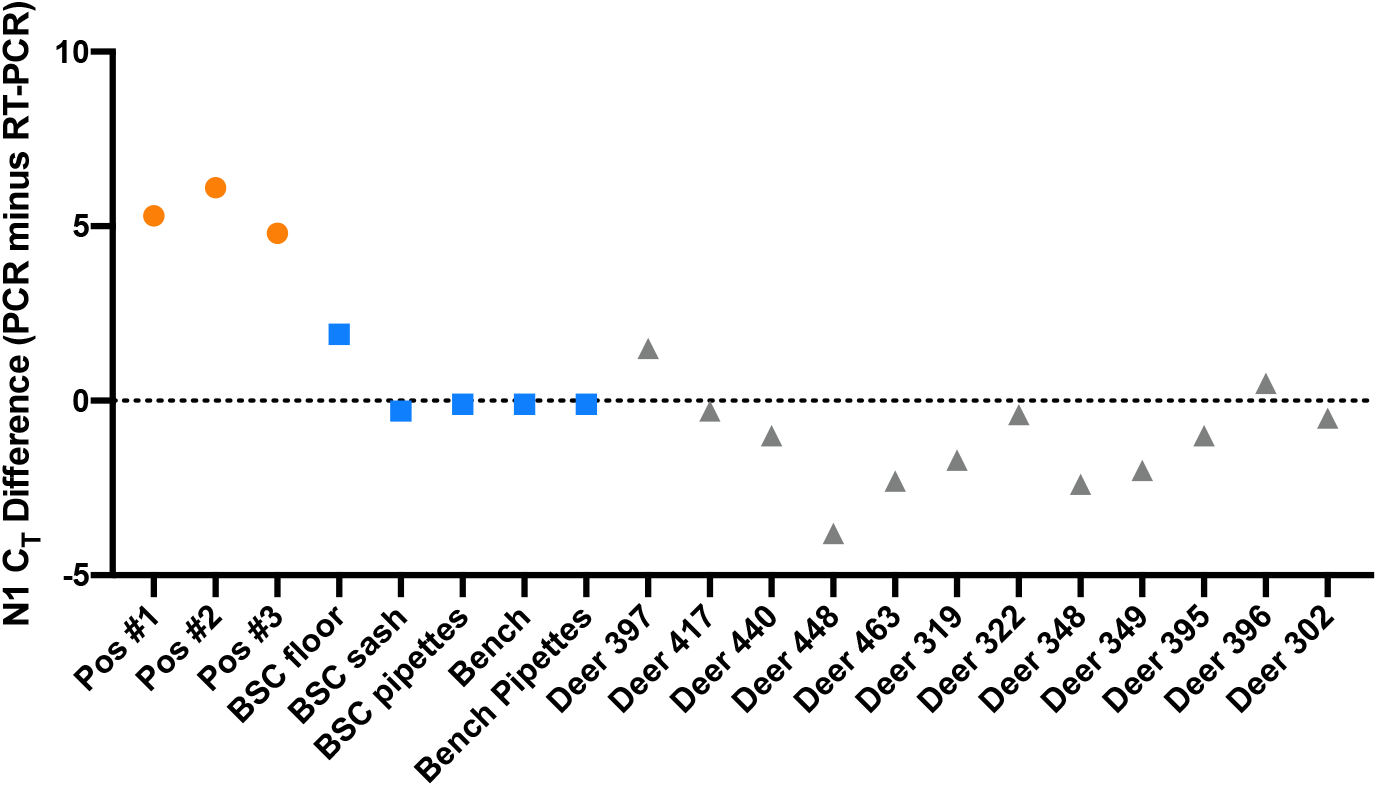
Environmental Swabs Show Contamination of SARS-CoV-2 Nucleocapsid DNA in Laboratory Environment. Samples from residual clinical SARS-CoV-2 positive specimens (pos #1-3), laboratory surfaces and equipment used to process/aliquot field samples (BSC=Biosafety Cabinet) and a selection of deer that initially tested positive for SARS-CoV-2 nucleic acid with the N1 primer set were analyzed by RT-PCR (to detect either RNA or DNA) and PCR (to detect RNA). The difference in cycle threshold (C_T_) between PCR and RT-PCR for each sample is shown. Positive control clinical samples shown in orange circles, environmental swabs shown in blue squares, deer samples shown in gray triangles.

Next, we compared qPCR and RT-qPCR amplification on select deer specimens that showed amplification with either the N1 or N2 primer sets (see Figure 2 for a subset and Dataset S2 for complete data). In each case, we were still able to detect viral nucleic acids, and as seen in the surface swabs the C_T_s were consistent between RT-qPCR and qPCR reactions (average N1 C_T_ 1.1±1.4 cycles lower in qPCR experiment than in RT-qPCR).

This result indicated that the original N1/N2 results were most likely detecting DNA contamination. The contamination likely occurred during the aliquoting step (after specimen collection), and illustrates the great difficulty posed by performing RT-qPCR based surveillance efforts in tandem with experiments that require the handling of plasmid DNA or PCR product without a physically separate facility, as previously reported^21–24^. Given the similar C_T_ values for both the RT-qPCR and qPCR of laboratory surfaces and deer specimen samples, as well as the presence of N gene plasmid DNA (and associated contamination) in the laboratory where the samples were aliquoted, we concluded that the 2022 N1/N2 results were false positives.

To accurately detect the presence of SARS-CoV-2 viral RNA in the 2022 season samples, we repeated our RT-qPCR analysis using a new and independent set of primers, this time targeting the E gene^25^ rather than N. There was no E gene plasmid DNA present in the laboratory in which these samples were processed and aliquoted, and no E amplification products were present at any point in the study. All 474 samples from the 2022 season were undetectable by the E gene primer/probe set, indicating that there was no detectable SARS-CoV-2 viral RNA in any Vermont wildlife surveilled during the 2021 or 2022 seasons (Dataset S1).

## Discussion

White-tailed deer can both successfully be infected with and transmit SARS-CoV-2. This has been demonstrated by both laboratory studies^16,17^ and several reports of naturally infected deer in multiple states and provinces within the United States and Canada^4,9–15^. Since prior surveillance studies have reported RT-qPCR positivity rates of upwards of 30% in nasal swabs^4^ and seropositivity rates of more than 40%^12,26^, it was initially surprising that no animals within the Vermont sample set were positive, especially during the 2022 season. However, recent work from Diel et al. describing the spread of SARS-CoV-2 within deer in New York state during the 2021 and 2022 seasons showed only sporadic positives during 2021 and a significant increase (up to 20%) in the 2022 season^15^. Furthermore, the majority of positive cases were detected in the western half of New York and near New York City, the farthest regions geographically from the Vermont border^14^. A second study furthers this argument, revealing the relatively low positivity rate of 1.2% within the Quebec province in Canada, directly north of Vermont^27^. Therefore, SARS-CoV-2 may be circulating in Vermont deer at a low level. To assess our ability to detect this, we performed a power analysis to calculate the probability of detecting one case of SARS-CoV-2 within our 472 samples from the 2022 deer season as a function of underlying SARS-CoV-2 prevalence. If we were to repeat our surveillance efforts, we would expect to find at least one positive sample 80% of the time if the underlying prevalence were at least 0.34%; similarly, we have 95% power to detect from a population that was 0.64% positive, and there is only a 1% chance of our sampling no positives if the population were 0.97% positive (Figure S2).

While it has not been established how SARS-CoV-2 is introduced into wild deer populations, it seems likely that this occurs via human-to-deer transmission, deer-to-deer transmission, or a combination of the two^26,28^. Vermont may have multiple features that reduced the risk of human-to-deer transmission so far in the COVID-19 pandemic. First, the state of Vermont is sparsely populated in general, but especially in many of the places where deer are hunted, therefore reducing the potential for human-deer interaction. Additionally, the number of COVID-19 human infections within the state of Vermont was much lower than most other places within the USA (including neighboring states with higher levels of SARS-CoV-2 detected in deer) during the period in which we were conducting surveillance (Figure 3, Dataset S3).

**Figure 3.**
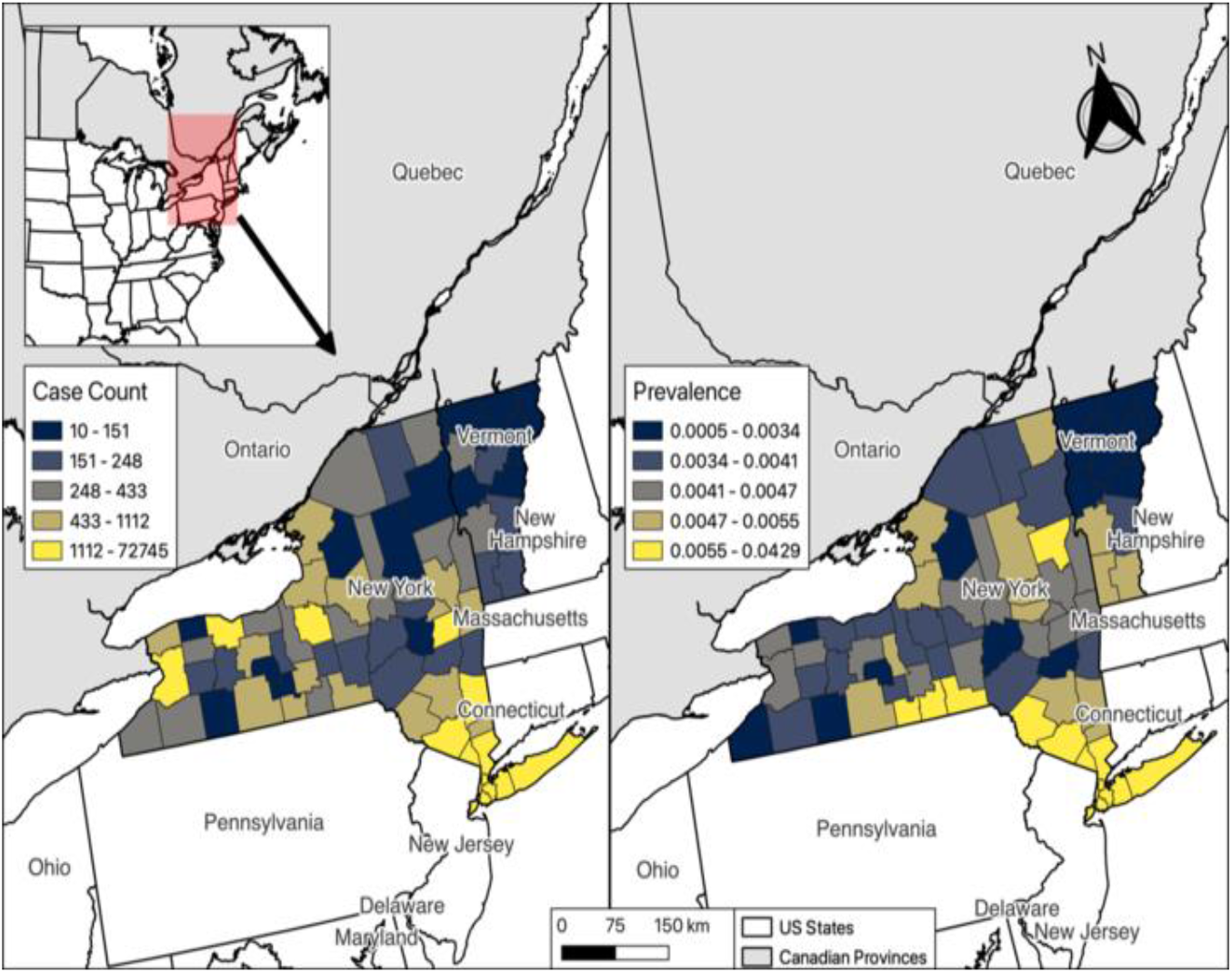
Case Counts and Prevalence of COVID-19 in Vermont and New York. Geographic distribution of COVID-19 cases (reported by the Vermont Department of Health and New York State Department of Health) during the during the surveillance period for the 2022 season (Oct 15^th^-Nov 15^th^ 2022) at the county level. Raw case counts are shown on left and prevalence (case counts/county population) is shown on right (population counts are an estimate based on US Census 2022 data). Graphs were generated using QGIS.

Finally, Vermont lacks an established deer farm industry, with only three farms reported in 2017^29^, all which contain cervid species other than white-tailed deer since it is illegal to have captive white-tailed deer in Vermont. This agricultural set-up decreases the number and duration of direct contact between humans and cervids in the state. Transmission between farmed animals (such as mink) and farm employees that care for them has been well documented for viruses including SARS-CoV-2 and is a plausible route for the initial introduction of SARS-CoV-2 into deer populations as well^30^. Texas, Pennsylvania, Indiana, Ohio, and Michigan alone account for over 65% of deer farms within the United States^29^ and several of these states have also reported high rates of SARS-CoV-2 prevalence in captive and/or wild deer^9,11–13,30^.

A limitation of this study is the lack of samples other than nasal swabs, such as retropharyngeal lymph nodes or blood samples. Retropharyngeal lymph nodes (RPLNs) are commonly collected as part of surveillance efforts for chronic wasting disease; however, VT only conducts surveillance for this disease when warranted by clinical signs/symptoms currently, and not on hunter harvested deer. A 2022 study from Ontario, Canada reported a 2.3% (5/213) positivity in nasal swabs, compared to a 6% (17/298) positivity in retropharyngeal lymph nodes within the white-tailed deer they sampled, potentially demonstrating the increased sensitivity of RPLNs samples to detect SARS-CoV-2 in this species^10^.Since no blood samples or lymph nodes were collected in this study, we were unable to perform serology experiments to detect the presence of SARS-CoV-2 antibodies that would reveal SARS-CoV-2 disease history. The results reported here only represent a lack of active infections in the animals surveilled at a single discrete timepoint. While information into the natural history of SARS-CoV-2 infections in wildlife during the 2022 season would be highly informative, the lack of standard collection of blood samples at Vermont hunting check stations was logistically prohibitive for the acquisition of these samples during the 2021-2022 hunting seasons.

While our findings that there does not appear to be widespread SARS-CoV-2 in Vermont deer are reassuring at present, we do not expect this to continue indefinitely considering the increasing cases detected in the wildlife of neighboring regions. Surveillance efforts to help detect the transmission and adaptation of SARS-CoV-2 in wildlife should be established throughout North America and should ideally prioritize species susceptible to infection. Ongoing surveillance studies will be required to understand not only the status of SARS-CoV-2 in Vermont wildlife populations, but also to understand the transmission and spread of the disease over time. Efforts to monitor the prevalence and mutational changes in SARS-CoV-2 viral genome are especially important within common and social species, such as white-tailed deer. The human health implication of deer as a SARS-CoV-2 reservoir is a sincere concern and warrants continued surveillance as a crucial measure in pandemic preparedness.

## Methods

### Wildlife Specimen Procurement

All samples were collected in collaboration with the Vermont Agency of Natural Resources, Department of Fish & Wildlife. All white-tailed deer and bear samples were collected during the Vermont hunting season. For the 2021 season, deer samples were collected on the opening weekend of rifle season (November 12^th^, 2022). For the 2022 season, samples were collected on youth weekend (Oct 22-23^rd^, 2022) and the opening weekend of rifle season (November 12^th^-13^th^, 2022). During these dates, samples were collected across the state of Vermont from deceased animals brought to big game reporting stations by hunters.

For fur-bearing animals (i.e., foxes, otters, coyotes, bobcats, and fishers), whole-animal carcass specimens were collected throughout the entirety of 2021 and stored at -20°C until SARS-CoV-2 swab sample collection occurred in a batch-wise fashion during February-March 2022. Most whole-animal specimens were collected between October 2021 and March 2022 and therefore stored for only a few months (details for individual specimens available in Dataset S1).

Nasal swabs were collected by inserting a dry, sterile swab (Copan #164KS01) approximately 1 inch into each nasal cavity of the specimen and making five passes around the interior of the nostril, ensuring even contact with the wall of the cavity. If the nasal cavity was inaccessible, throat swabs were taken by inserting the swab as far back into the throat as possible and making five passes around the entire circumference (denoted in Dataset S1).

Samples were stored in 3mL of phosphate-buffered saline (Gibco #10010023) on ice until returning to the laboratory where they were transferred to -80°C until further use.

### Environmental Swabbing for Laboratory Plasmid Contamination

Environmental contamination samples were collected by rubbing the surface of interest with a dry, sterile swab (Copan #164KS01) for approximately 10 seconds, rolling the swab during this time to ensure maximal surface contact. Samples were stored in 1mL of phosphate-buffered saline (Gibco #10010023) and stored at -80°C until nucleic acid extraction and amplification could occur.

### Nucleic Acid Extraction & Amplification

All 2022 season samples were thawed once and aliquoted at the University of Vermont between sample collection and extraction. All further processing and testing of swabs took place in a Clinical Laboratory Improvement Amendments (CLIA)- and College of American Pathologists (CAP)-certified facility at the University of Washington Virology Laboratory.

Total nucleic acids (TNA) were extracted using Roche MagNA Pure 96 instruments as previously described^31^, with 200µL of swab liquid extracted and eluted into 50µL. Each extraction plate included a positive control (pooled SARS-CoV-2-positive clinical remnants) and a negative control (cells derived from a HeLa cell line). All amplifications used AgPath ID One-Step RT-PCR enzyme and master mix (Life Technologies, ThermoFisher, Cat. #4387424M) and 10µL of TNA per reaction and were carried out on ABI 7500 thermocyclers. In addition to the positive and negative controls from each extraction, each amplification plate contained a No-Template negative control (NTC; water). One of two primer/probe sets was used in all reactions: WHO-E^25^; or multiplexed CDC N1 and N2^32^. RT-qPCR amplifications consisted of 10’ at 48° C (reverse transcription), 10’ at 95° C (Reverse Transcriptase inactivation / polymerase hot-start), and 40 cycles of 15” at 95° C and 45” at 60° C. qPCR amplifications used the same cycling conditions but omitted the initial 10’ at 48° C step. EXO RNA was added to all samples prior to extraction, and EXO amplification was included in every RT-qPCR reaction as an internal control to monitor for RNA degradation and PCR inhibition^32^.

### Data Availability

All code, supplemental manuscript metadata, and supporting information can be found in GitHub @emilybrucelab (https://github.com/emilybrucelab).

## Acknowledgements

This study was supported by NIH grant P30GM118228-04 and UVM start-up funds (EAB) and an American Heart Association predoctoral fellowship (HWD). We would like to acknowledge the Vermont Agency of Natural Resources, Fish & Wildlife Department for allowing us to participate in all their ongoing research efforts to collect these samples. We would also like to thank the students of the Wildlife & Fisheries Club at the University of Vermont for helping collect samples. Finally, a big thank you to all the Vermont hunters that were willing to participate in our efforts.

## Supplemental Figures

**Supplemental Figure 1.**
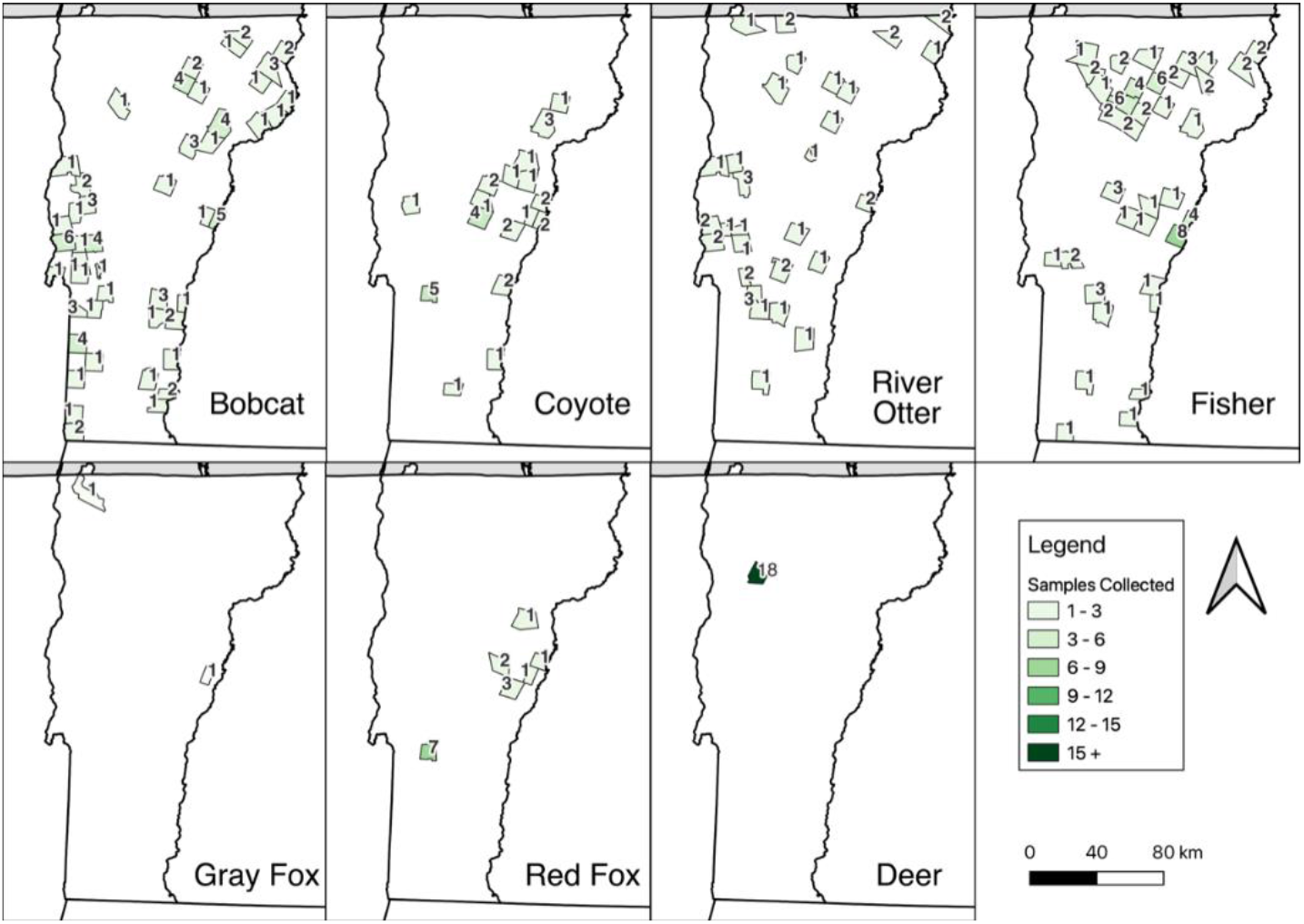
Geographic distribution of 2021 sample collection by species. Graphs generated using QGIS.

**Supplemental Figure 2:**
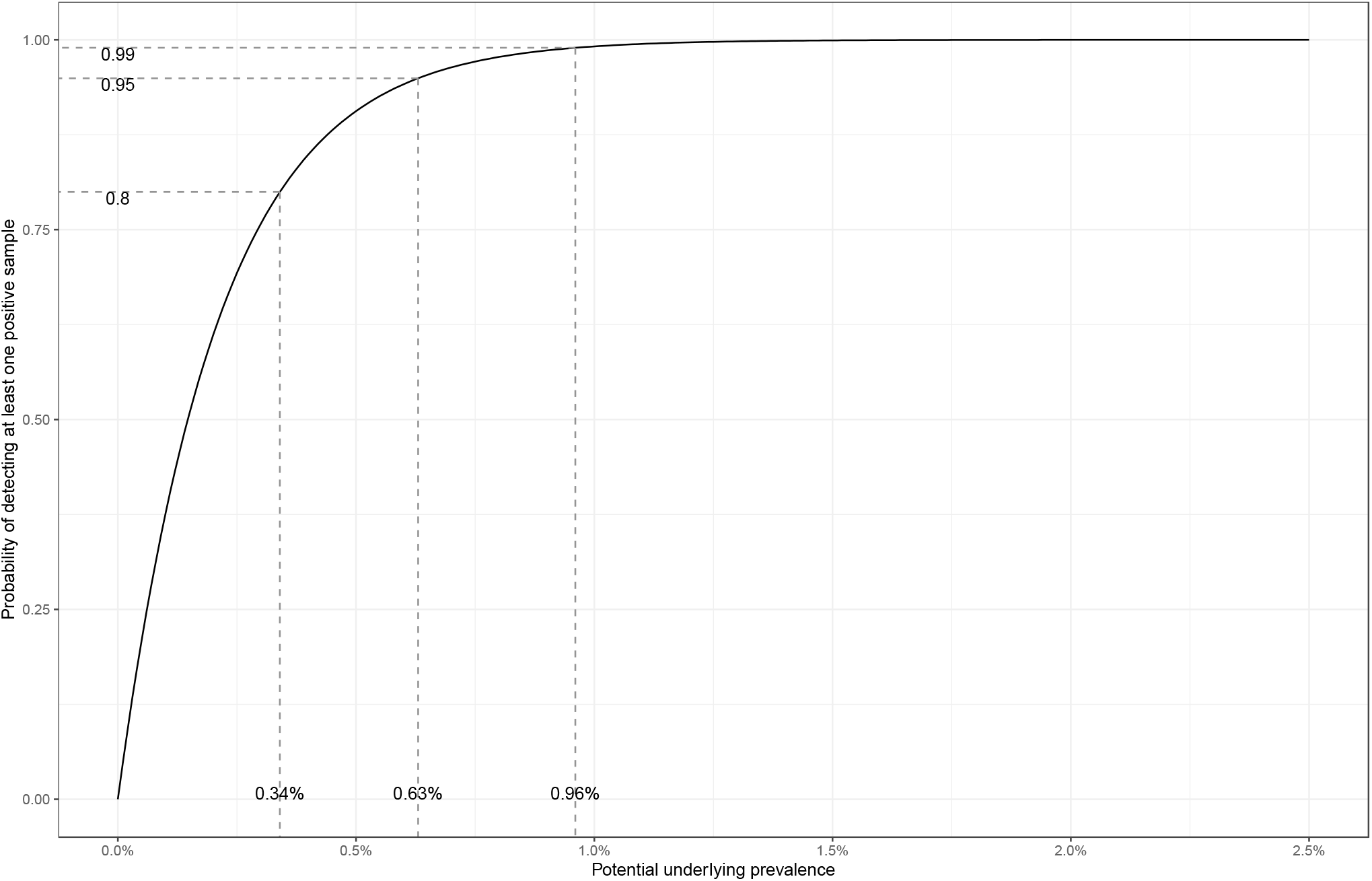
Power to Detect a Given Prevalence. Using the binomial distribution for 472 trials (the number of deer samples collected in the 2022 hunting season), we calculated the probability of at least one success (SARS-CoV-2 detection) as a function of the unknown underlying SARS-CoV-2 prevalence (percent positivity).

